# Investigating the Impact of Microplastics on Fish Muscle Cell Proliferation and Differentiation: Enhancing Food Safety in Cultivated Meat Production

**DOI:** 10.1101/2023.10.11.561915

**Authors:** Taozhu Sun, Alfonso Timoneda, Amiti Banavar, Reza Ovissipour

## Abstract

Cultivated meat, a sustainable alternative to traditional livestock farming, has gained attention for its potential environmental and health benefits. However, concerns about microplastic contamination pose challenges, especially when sourcing cells from marine organisms prone to microplastic bioaccumulation. Additionally, the pervasive presence of microplastics in laboratory settings, ingredients, and during the production, increases the risk of unintentional contamination. This study focused on Atlantic mackerel (*Scomber scombrus*) skeletal muscle cell lines to examine the effects of microplastic exposure, represented by fluorescent polyethylene microspheres (10-45 µm) on cell performance including cell proliferation, cell viability, gene expression, and differentiation processes critical for cultivated meat production. The results revealed significant impacts on cell attachment and proliferation at microplastic concentrations of 1 µg/mL, 10 µg/mL, and 50 µg/mL. Notably, the 10 µg/mL concentration exerted the most pronounced effects on cell viability during both attachment and proliferation phases. While the results indicated that both microplastic concentration and size influence cell viability, cell differentiation remained unaffected, and additional contributing factors require further investigation. These findings underscore the necessity of thoroughly exploring microplastic-cell interactions to ensure food safety and safeguard health within the burgeoning cultivated meat industry.

## 1. Introduction

Cultivated meat, derived from the cultivation of animal cells, presents an innovative shift in food production with potential environmental and health benefits (Dupuis et al., 2023; Eibl et al., 2021; Jahir et al., 2023; Rischer et al., 2020). Produced within controlled environments, this approach not only minimizes the risks associated with conventional farming contaminants but also promises a more resource-efficient methodology (Stephens et al., 2018). Recent studies indicate that, with renewable energy integration, cultivated meat could achieve up reductions of global warming by 92%, air pollution by 93%, land use by 95%, and water consumption by 78% compared to traditional beef farming (Kim et al., 2022; Sinke et al., 2023; Vergeer et al., 2021). As the industry advances, cultivated meat is projected to command a substantial portion of the $1.7 trillion conventional meat and seafood market, addressing pressing challenges like deforestation, biodiversity loss, and antibiotic resistance (Sinke et al., 2023; Vergeer et al., 2021).

In controlled laboratory environments, cultivated meat is produced from cells, such as those from animals. These cells undergo proliferation in specialized growth mediums to form muscle tissue, representing a potentially safer, more ethical, and environmentally sustainable alternative to conventional meat production (Chriki & Hocquette, 2020; Ong et al., 2021). However, a potential safety concern in this innovation is the contamination of microplastics. One avenue of potential contamination arises from the source animals. Marine ecosystems, for instance, are known reservoirs of microplastics (Andrady, 2011; Cole et al., 2011; Ivar do Sul & Costa, 2014). This results in bioaccumulation within marine life, such as fish and oysters (Bhuyan, 2022; Courtene-Jones et al., 2022; Galloway et al., 2017; Sharma & Chatterjee, 2017). When such marine organisms serve as the source animals for cell extraction, undetected microplastics could be inadvertently introduced into the cultivation process. Existing analytical methodologies often fail to detect smaller microplastic particles, leading to potential underestimations of their abundance in source organisms (Adhikari et al., 2022; Huppertsberg & Knepper, 2018; Lv et al., 2021; Vivekanand et al., 2021). Another significant source of contamination is the laboratory environment itself. Studies have underscored the pervasive nature of microplastics in laboratory settings, emanating from the degradation of ubiquitous plastic equipment, containers, and consumables (Koelmans et al., 2019; Löder et al., 2017; Schymanski et al., 2018). The production process of cultivated meat necessitates the use of various plastic-based apparatus, including bioreactors, pipettes, cell culture flasks, and other equipment that come in direct contact with the medium and growing cells (Allan et al., 2019; Lee et al., 2022).

Microplastics, tiny fragments of plastic less than 5 millimeters in size, have garnered significant attention due to their ubiquity in the environment and the potential risks they pose to human health (Lim, 2021; Lwanga et al., 2022; Osman et al., 2023). Upon ingested, these particles can traverse the gastrointestinal tract, and some evidence suggests that smaller micro- and nanoplastic particles may even penetrate tissues, entering the circulatory and lymphatic systems (Campanale et al., 2020; Fournier et al., 2023; Hirt & Body-Malapel, 2020; Jiang et al., 2020; Kannan & Vimalkumar, 2021; Li et al., 2023; Ramsperger et al., 2023; Yee et al., 2021). These fragments can act as carriers for various toxicants, including heavy metals, polycyclic aromatic hydrocarbons, and endocrine-disrupting chemicals (Abbasi et al., 2021; Amelia et al., 2021; Campanale et al., 2020; Karla Lizzeth et al., 2023; Yee et al., 2021), thereby introducing these harmful agents into the human body. From a cellular perspective, the risks of microplastics become more intricate. The direct interaction between cells and microplastics can lead to physical disruptions, such as membrane damage (Dai et al., 2022; Fleury & Baulin, 2021; Wang et al., 2022), and chemicals inherent to or leached from these plastics are known to induce oxidative stress, inflammatory responses, and genotoxic effects (Alqahtani et al., 2023; Cao et al., 2023; Goodman et al., 2021; Herrala et al., 2023; Hirt & Body-Malapel, 2020; Jeyavani et al., 2023; Mattioda et al., 2023). Potential risks of such interactions encompass DNA lesions, organ dysfunctions, metabolic irregularities, immunological aberrations, neurotoxicity, and perturbations in reproductive and developmental processes (Galluzzi et al., 2018). Furthermore, previous research has indicated a potential link between microplastic exposure and the development or exacerbation of certain chronic diseases (Lee et al., 2023; Wu et al., 2023). Given the documented adverse effects of microplastics upon ingestion, understanding and mitigating these risks is paramount for the cultivated meat industry (Chain, 2016; Mamun et al., 2023; Rubio-Armendáriz et al., 2022; Ziani et al., 2023).

While the presence and potential hazards of microplastics are increasingly acknowledged, understanding the exact mechanisms by which they influence cellular functions remains a critical research frontier (O’Neill & Lawler, 2021; Thornton Hampton et al., 2022). To understand the cellular impacts of microplastic exposure more comprehensively, we utilized Atlantic mackerel (*Scomber scombrus*) skeletal muscle cell lines, previously established and characterized by Saad et al. (2023), given their relevance to cultivated meat processing. This study employed fluorescent polyethylene microspheres (10-45 µm) as representative microplastics, a size range previously documented in fish (Makhdoumi et al., 2023; Thiele et al., 2021). We aimed to elucidate the effects of microplastics on cellular performance, emphasizing cell viability during the attachment and growth phases, as well as cell differentiation, which are pivotal processes in cultivated meat production (O’Neill et al., 2021; Reiss et al., 2021). The study employed microplastic concentrations of 1 µg/mL, 10 µg/mL, and 50 µg/mL. Preliminary results indicated that all treatments significantly affected cell attachment (on day 2) and proliferation (on day 4), with no discernible effects on cell differentiation after two weeks. Variables such as microplastic size and concentration potentially influenced these outcomes. These findings, albeit initial, provide foundational insights for subsequent research, emphasizing the importance of understanding microplastic-cell interactions for ensuring food safety, protecting human health, and mitigating environmental impacts.

## 2. Materials and Methods

### 2.1. Cell lines preparation and maintenance

The mackerel cell lines (MACK2) used in this study were obtained from Dr. David L. Kaplan Lab at Tufts University. The cell preparation followed the protocol described in Saad et al. (2023). Briefly, frozen cells at passage 81 were thawed using 9 mL of complete growth medium, which comprised Leivovitz’s L-15 medium (Gibco™, Billings, MT, US) and supplemented with 20% fetal bovine serum (FBS, Gibco™, Billings, MT, US), 1 ng/mL FGF2-G3 (human) growth factor (Defined Bioscience, San Diego, CA, US), 20 mM HEPES (Gibco™, Billings, MT, US), and 1% Antibiotic–Antimycotic (Gibco™, Billings, MT, US). The cell suspension was then centrifuged at 500 RCF for 6 minutes, and the resulting pellet was resuspended in 10 mL of growth medium. The cells were incubated in a 75Lcm^2^ culture flask (Thermo Fisher Scientific, Waltham, MA, US) at 27 °C in a CO_2_ free incubator. Maintenance of the cells involved regular passaging at approximately 70% confluency and seeding at a density of approximately 5000 cells/cm^2^. Alternatively, cells were stored by freezing in growth medium supplemented with 10% dimethyl sulfoxide (DMSO, Sigma Aldrich, St. Louis, MO, US).

### 2.2. Microplastic preparation and exposure

Fluorescent green polyethylene microspheres obtained from *Cospheric LLC* (Goleta, CA, US) were used in this study. The microspheres exhibited a size range of 10-45 µm. Prior to experiments, a sterilization process was undertaken using 91% isopropyl alcohol (IPA), allowing excess fluid to drain as the spheres gradually underwent evaporation. Subsequently, the sterilized microspheres were integrated into the completed growth medium supplemented with 0.01% Tween 20. Microspheres were introduced into the experimental setup at concentrations of 1, 10, and 50 μg/mL based on previous studies (Hwang et al., 2020; Palaniappan et al., 2022; Schirinzi et al., 2017). An experimental control consisting of the complete growth medium with 0.01% Tween 20 but devoid of microspheres was included in the study. To achieve a consistent distribution of the microspheres within the growth medium, a pre-experimental step involved subjecting the microsphere medium to sonication prior to each experiment.

### 2.3. Effects of microplastic on cell attachment and viability

Mackerel cells were seeded in triplicate in the 6-well plates at a seeding density of approximately 5000 cells/cm². The cells were cultured at a constant temperature of 27°C and devoid of CO^2^. To investigate the influence of microplastics on different stages of cell growth, two distinct experimental conditions were employed. In the first scenario, microplastics were incorporated into the cell medium prior to seeding to assess their influence on cell attachment, a crucial initial step in cell proliferation. After 48 hours of incubation, cells reached the logarithmic growth phase and were detached from the plate surface using 0.25% trypsin–EDTA (Thermo Fisher Scientific, Waltham, MA, US). The cell viability was evaluated using the Trypan Blue Assay (Strober, 2019) and the Countess 3 FL Automated Cell Counter (Invitrogen™, Thermo Fisher Scientific, Waltham, MA, US). In the second scenario, microplastics were introduced to the cell medium after the logarithmic growth phase was attained, with the old medium replaced by either microplastic-containing or control fresh medium. Subsequent to the introduction of microplastics, the cells remained undisturbed for a period of 4 days, allowing for the exploration of potential interactions between microplastics and the cells. After this interaction period, the cells were detached, and the viable cell count was determined, offering insights into the effect of microplastics on cell growth after the initial proliferation stages.

### 2.4. Effects of microplastic on cell differentiation

#### 2.4.1. RT-qPCR gene expression analysis

Mackerel cells at passage 82 were detached from 6-well flasks using trypsin for 3-4 min and pelleted by a 7.5 min centrifugation at 500 RCF. RNA was extracted from samples using the NucleoSpin RNA kit (Mackerey-Nagel, Dueren, Germany) and quantified with a Qubit 4 Fluorometer using the Qubit RNA High Sensitivity (HS) assay kit (Thermo Fisher Scientific, Waltham, MA, USA). cDNA libraries for each sample were constructed from 100 ng of RNA using the PrimeScript RT master mix (Takara Bio, Kutatsu, Japan) under manufacturer specifications. A minus reverse transcriptase control (-RT) was made from the sample exhibiting the higher RNA yield by not adding the master mix to the sample. The PrimeScript RT master mix contains both random hexamers and oligo dT primers. RT-qPCR was performed in a CFX Opus 96 thermocycler (Bio-Rad, Hercules, California, USA) using the TB Green Advantage qPCR premix (Takara Bio, Kutatsu, Japan) and the oligonucleotides specified in Table 1. Primers were designed using a reference genome for southern bluefin tuna (Thunnus maccoyii; NBCI RefSeq GCF_910596095.1) by Saad et al. (2023). Three technical replicates were performed per sample and gene, as well as for the - RT and no template controls (NTC). Amplification conditions were as follow: initial step of 30 s at 95°C, followed by 40 cycles of 5 s at 95°C and 30 s at 60°C, and a final dissociation analysis of 15 s at 65°C and 0.5°C/s increases until reaching 95°C. Absolute gene expression values were calculated as 2^(-ΔCt) using the hypoxanthine guanine phosphoribosyltransferase (HPRT) gene as the housekeeping gene. Relative gene expression values were calculated as 2^(-ΔΔCt) over the 0 ng/mL microplastics control.

**Table 1.**
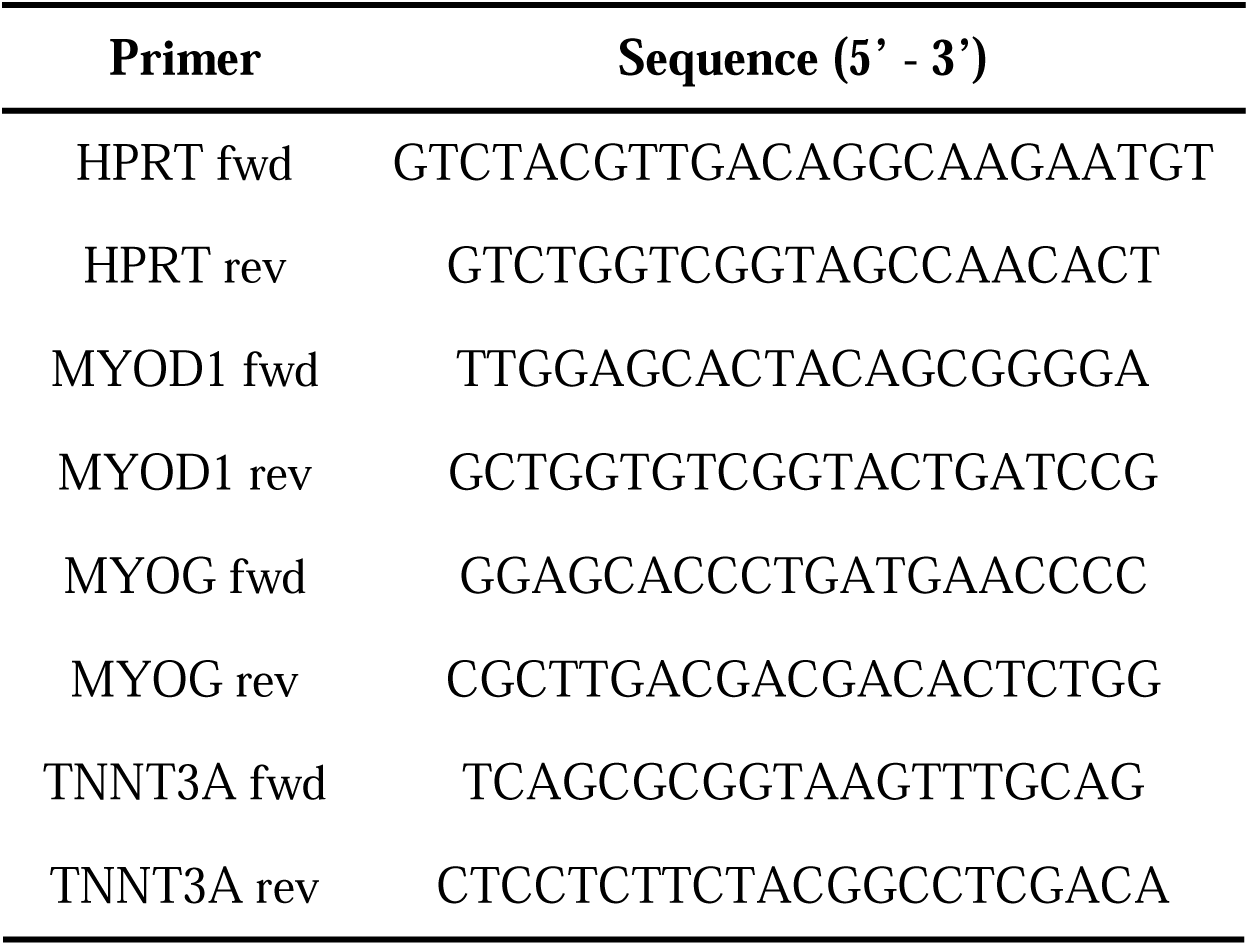
Oligonucleotides used for RT-qPCR of mackerel cells.

#### 2.4.2. Immunostaining

To observe the influence of microplastics on cell differentiation, immunostaining for myosin heavy chain was conducted following the protocol outlined in Saad et al. (2023). Mackerel cells were cultured at 100% confluency in growth medium and exposed to microplastics at varying concentrations, with subsequent observation over a 14-day differentiation period. Following culture, cells were fixed with 4% paraformaldehyde at room temperature for 30 minutes (Thermo Fisher Scientific, Waltham, MA, USA). Subsequently, the cells were rinsed with Phosphate Buffered Saline (PBS, Sigma Aldrich, Burlington, MA, USA) and permeabilized for 10 minutes using 0.1% Triton-X (Sigma Aldrich, Burlington, MA, USA). Subsequent to permeabilization, the cells were blocked for 30 minutes using 1× blocking buffer (Abcam, Cambridge, UK), followed by an additional PBS wash. The primary antibody solution, MF-20 (4 μg/mL), was applied to the cells and allowed to incubate overnight at 4°C. After a subsequent PBS wash, the cells underwent an additional 30-minute blocking step using 1× blocking buffer and were then incubated for 1 hour with secondary antibodies – Goat Anti-Mouse IgG H&L (Alexa Fluor® 594, Abcam, Cambridge, UK), and Phalloidin-iFluor 488 Reagent (Abcam, Cambridge, UK) – each diluted at 1:1000 in 1× blocking buffer. After a final wash with PBS, the cell nuclei were stained with 4′,6-diamidino-2-phenylindole (DAPI, 1 μg/mL, Thermo Fisher Scientific, Waltham, MA, USA) in PBS for 15 minutes at room temperature. Imaging was conducted using a fluorescence microscope (DP27, Olympus Life Science, Tokyo, Japan) equipped with an LED Illumination system (CoolLED, Andover, UK).

### 2.5. Statistical analysis

Statistical analysis was conducted using One-way ANOVA, with a t-test utilized for comparisons between two parameters.

## 3. Results and Discussion

### 3.1. Cell viability

The viability of cells during both the attachment and growth phases is paramount to the cultivation of meat. In the attachment phase, cells must successfully anchor to a scaffold or matrix to prevent their loss during media changes, establishing a robust foundation for subsequent stages. Following successful attachment, it’s imperative for these cells to proliferate efficiently, ensuring an adequate cellular population for muscle tissue formation. Any significant cell mortality or diminished proliferation during these phases could undermine the entire production yield and efficiency (Allan et al., 2019; Bodiou et al., 2020). In order to investigate the effects of microplastics on mackerel cell viability at attachment and growth phases, microplastics were introduced to cells at different times in this study. Figure 1a highlighted the outcomes of microplastics on initial cell attachment when introduced into the cell medium before seeding; Figure 1b depicted the impact of microplastics on cell proliferation when added after the cells had reached the logarithmic growth phase. The initial number of cells for seeding was ∼ 5000 cells/cm², and the surface area of 6 well is 9.6 cm². The results were collected on Day 2 and Day 4 for Figure 1a and 1b, respectively, when large number of cells started to die and detached. In Figure 1a, the experimental control (no microplastic) reached to 410000; the cells for the ones with microplastics reduced to the half. In Figure 1b, the cells number for control was 110,0000; the ones with microplastics ranged from 510,000 to 730,000. In general, both sets of results exhibited significant effects of microplastics on cell viability compared to the experimental control. Remarkably, a conspicuous trend emerged, indicating that microplastic concentrations of 10 μg/mL elicited the most pronounced effects, surpassing the responses observed at 1 and 50 μg/mL concentrations.

**Figure 1.**
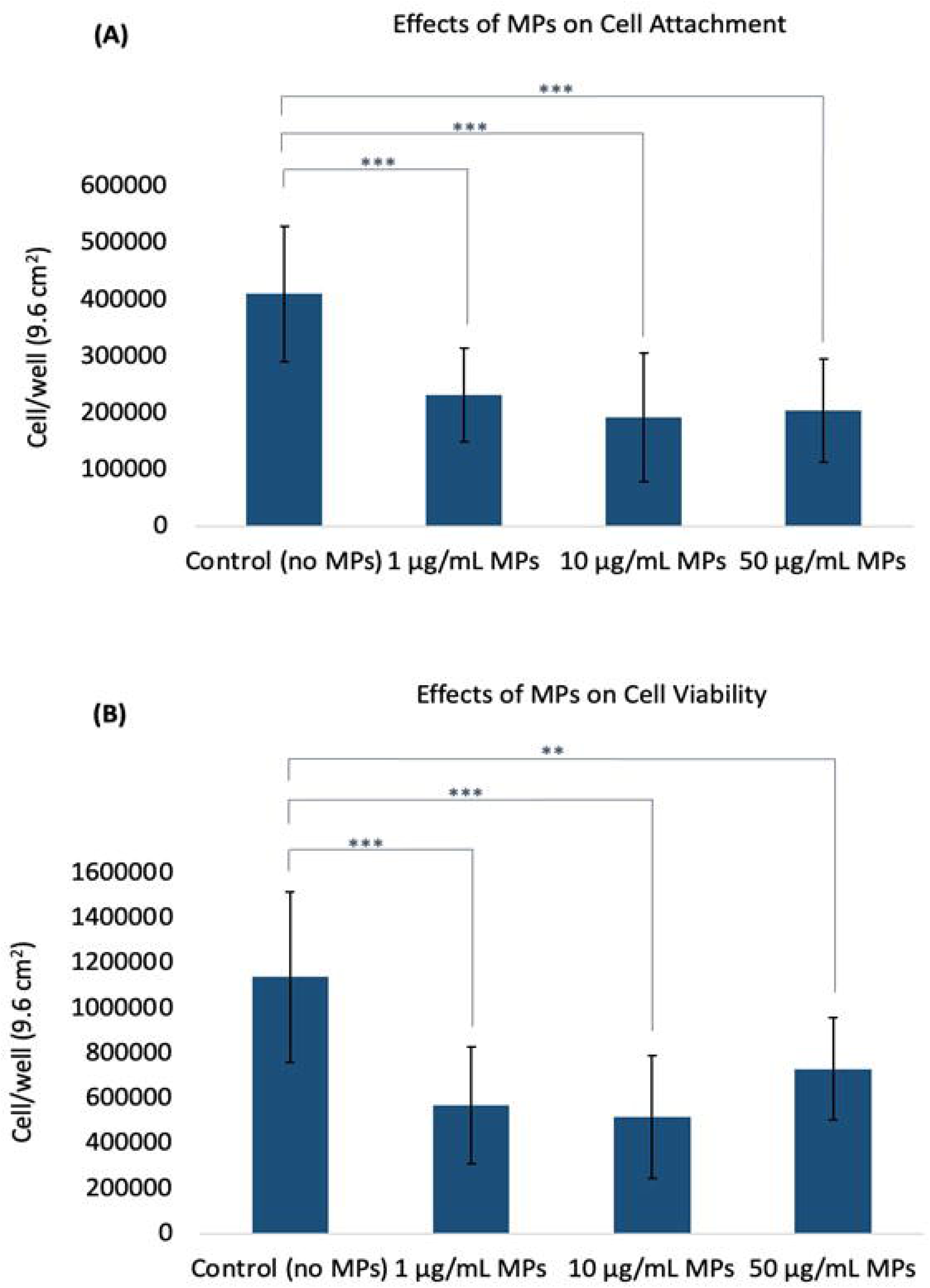
Effects of microplastics on cell viability. (A) Influence of microplastics on initial cell attachment when introduced to the cell medium prior to seeding. (B) Impact of microplastics on cell proliferation after cells reached the logarithmic growth phase. Error bars represent standard deviation. The t-test was used to evaluate the differences between each MPs treatment with experimental control; asterisks indicate a significant difference at P < 0.01 (**) and P < 0.001 (***).

#### 3.1.1 Concentration effects on cell viability

Our findings demonstrate a significant impact of microplastics (MPs) on mackerel cell viability, with the observed effects being potentially dose dependent. At tested concentrations of 1, 10, and 50 µg/mL, the 1 µg/mL dose demonstrated the lowest adverse effects, supporting the notion that MP toxicity is dose-dependent. This aligns with previous studies that observed similar trends in cell viability with MP exposure. For instance, Palaniappan et al. (2022) performed an investigation involving L929 murine fibroblasts and MDCK epithelial cell lines and noted a dose-dependent decrease in cell viability when exposed to 1, 10, or 20 μg/mL of PE or PS microspheres. Furthermore, their study highlighted amplified oxidative stress at higher MP doses, as evidenced by increased SOD3 gene expression. In another study (H.-S. Lee et al., 2021), human umbilical vein endothelial cells (HUVECs) were exposed to polystyrene microplastics (PS-MPs, 0 −100 μg/mL), revealing that higher doses markedly reduced cell viability and disrupted angiogenic tube formation in the short term, while inducing autophagic and necrotic cell death after prolonged exposure. Besides, in a distinct investigation focusing on the human intestinal milieu (Herrala et al., 2023), researchers assessed the toxicological ramifications of ultra-high molecular-weight polyethylene particles (250 −1000 μg/ mL) on human colorectal adenocarcinoma Caco-2 and HT-29 cells. A 48-hour exposure to these polyethylene particles precipitated a dose-dependent decline in cell viability and a concomitant upsurge in oxidative stress. The oxidative damage was particularly pronounced in the mitochondria, illuminating the broader health concerns.

The intricate interplay between MP concentration and its potential cytotoxic effects is evident. While a substantial proportion of literature supports a dose-dependent decrease in cell viability, certain exceptions persist. In a recent in vitro study examining the impact of microplastics (PVC and PE) on gilthead seabream and European sea bass head-kidney leucocytes (HKLs)(Espinosa et al., 2018), it was observed that exposure to varying concentrations of microplastics for 1 and 24 hours did not significantly affect HKL cell viability. Additionally, high doses of microplastics resulted in minimal changes to key cellular innate immune functions, including a decrease in phagocytosis and an elevation in respiratory burst activity. These divergent findings underline the significance of further investigations, accounting for microplastic type, size, and the specific cellular environment, to draw conclusive inferences on the broader impacts of MPs on cellular health.

#### 3.1.2 Influence of microplastic size and aggregation

In addition to concentration, particle size emerges as a critical determinant of microplastic (MP) toxicity. Our surprising observation, where the 10 µg/mL concentration exhibited more pronounced adverse effects compared to the ostensibly higher 50 µg/mL concentration, suggests that MP-induced effects might not simply be linear with concentration. Our hypothesis posits that this counterintuitive trend could be attributed to larger aggregates formed in the 50 µg/mL medium (as seen in Figure 2), which might mitigate direct cellular interactions and thus lessen their effects on cell viability. A systematic review assessed dose– response relationships regarding microplastics (MPs) and cell viability by evaluating studies up to March 2021(Danopoulos et al., 2022). Of the 17 studies reviewed, 8 were included in a meta-regression analysis. The review identified four MP-associated effects: cytotoxicity, immune response, oxidative stress, and barrier attributes, with genotoxicity showing no effect. Key predictors of cell death were irregular MP shape, exposure duration, and MP concentration. Notably, Caco-2 cells displayed heightened susceptibility to MPs. Concentrations as low as 10 μg/mL (5–200 µm) affected cell viability, while 20 μg/mL (0.4 µm) influenced cytokine release. These findings are consistent with our observations, particularly the pronounced effects we observed at 10 µg/mL MPs on cell viability.

**Figure 2.**
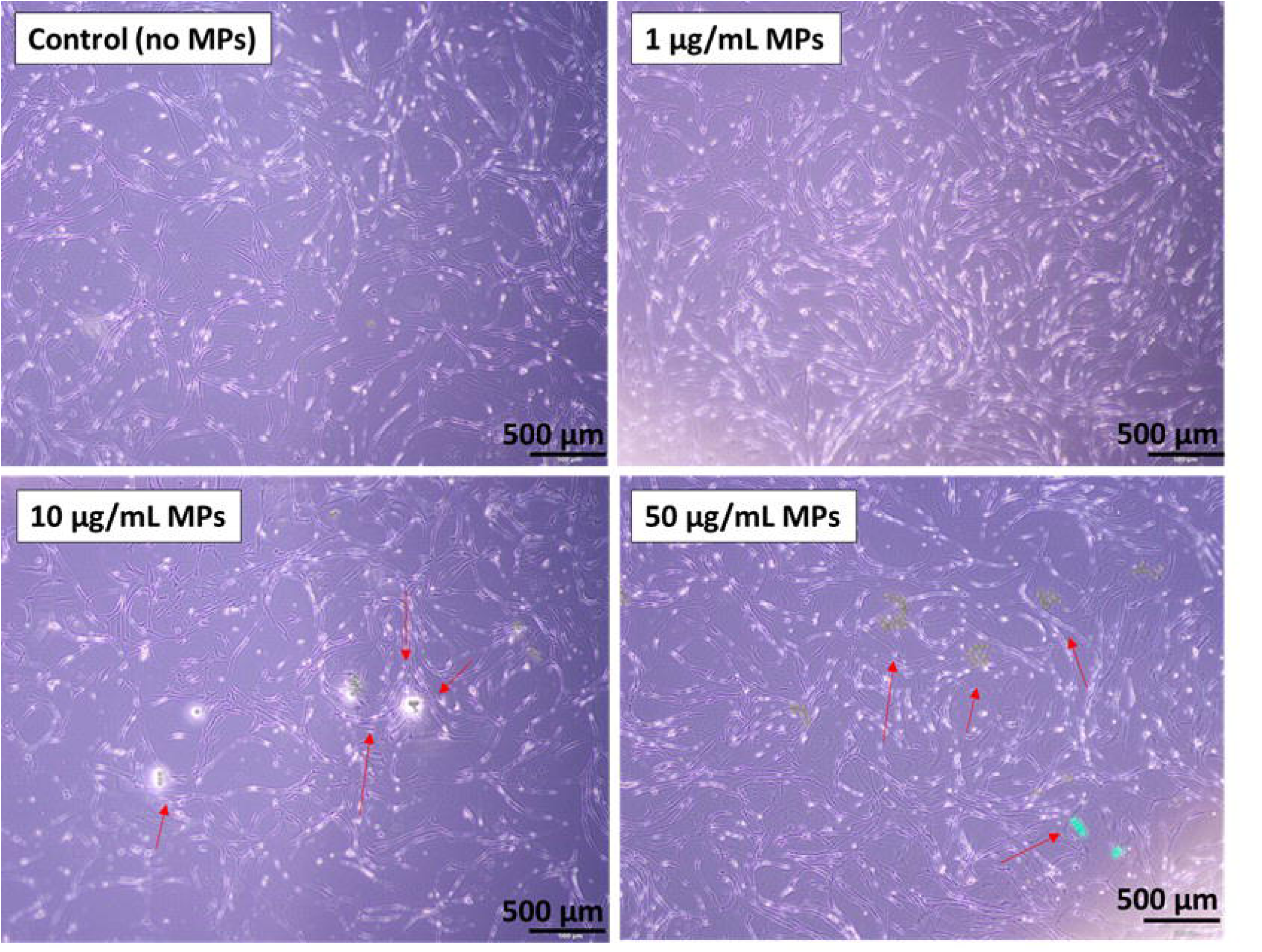
Morphological observations of mackerel cells post incubation with microplastics. Representative images of mackerel cells following a 4-day incubation, with and without microplastics (MPs). Microplastic aggregates are indicated by red arrows.

In a detailed investigation by Zhang et al. (2022), the impacts of polystyrene microspheres (PS-MPs) and nanospheres (PS-NPs) were explored across four distinct sizes: 0.1, 0.5, 1, and 5 μm. This research identified a marked preference for cellular uptake among the smaller nanoparticles compared to their larger counterparts. Notably, the PS-MPs presented minimal effects on cell viability and apoptosis. However, subtle indications of oxidative stress were discernible in high-concentration groups. A striking differentiation was evident in membrane damage, with PS-MPs inducing substantially greater damage than PS-NPs, emphasizing the size-dependent cellular responses to MPs. Building on the effects of microplastic size on cells, an investigation into polystyrene (PS) particle toxicity further elucidated these intricate relationships (Hwang et al., 2020). Researchers found that PS particles act as potential immune stimulants, triggering cytokine and chemokine production in a manner determined by size and concentration. Larger PS particles (10-100 µm in diameter) exhibited negligible cytotoxicity. In contrast, smaller particles, specifically those at 460 nm and 1 µm, detrimentally affected red blood cells. Their increased surface area, which facilitates stronger interactions like van der Waals forces, was pinpointed as the cause for hemolysis. Furthermore, exposure to these smaller PS particles resulted in elevated IL-6 secretion, signifying potential early-stage inflammation. However, the study also highlighted no substantial rise in histamine secretion, mitigating concerns of histamine-driven inflammation or allergic reactions. Particle uptake predominantly occurred through endocytosis and phagocytosis by phagocytic cells, leading to localized inflammation via pro-inflammatory cytokine release, rather than inducing direct cytotoxicity (Hwang et al., 2020). In addition to the examples described, other studies have consistently indicated that the size of microplastic particles significantly influences cellular interactions. Notably, smaller particles are linked to heightened cellular uptake, more pronounced inflammatory responses, increased apoptosis rates, and enhanced cellular stress responses (Revel et al., 2018; Wright & Kelly, 2017; Yong et al., 2020). These findings highlight the potential health risks associated with finer microplastic particles.

#### 3.1.3. Potential interactions and other involved factors

The interaction between cells and microplastics is a multifaceted process, influenced by a confluence of factors, such as the physicochemical properties of microplastics, cellular characteristics, and prevailing environmental conditions (Campanale et al., 2020; Revel et al., 2018; Smith et al., 2018). This dynamic interplay gains significance when contemplating the potential ramifications of microplastics on cellular health (Lee et al., 2023; Li et al., 2023). While this study predominantly illustrated the influence of microplastic size and dose on cell viability, it’s vital to embed these insights within the expansive framework of factors modulating microplastic-cell interactions. The polymer composition (e.g., polymer type, presence of additives, and potential for the microplastics to absorb other environmental contaminants) of microplastics, for instance, is often associated with discrete cytotoxic effects (Duis & Coors, 2016; Hwang et al., 2020; Revel et al., 2018). The shape of microplastics further refines this interaction spectrum. Fibrous microplastics have been reported to potentially introduce physical harm, further affect tissues or induce blockages (Watts et al., 2015; Wright et al., 2013), whereas spherical microbeads, prevalent in personal care products, could facilitate a smoother cellular internalization(Wright et al., 2013).

Nanoscale microplastics can lead to the creation of Reactive Oxygen Species (ROS), suggesting that they may cause stress to cells by promoting oxidative reactions (Campanale et al., 2020; Paul et al., 2020; Yee et al., 2021). Concurrent inflammatory responses may destabilize cellular homeostasis, possibly marking the onset of apoptosis (Elmore, 2007; Lamichhane et al., 2023; Wright et al., 2013). There could also be direct physical effects, such as potential damage or blockages in tissues, especially in organisms with multiple cell types (Bhagat et al., 2021; Yee et al., 2021). Certain cellular stress signals, specifically p-JNK and p-p38, emphasize that microplastics can be seen as stress-causing agents (Jeong et al., 2017; Jeong et al., 2016; Scopetani et al., 2020). It’s also worth noting that different organisms and cell types can react differently to these plastics (Bhagat et al., 2021; Jeong & Choi, 2019). Comprehending these multifaceted interactions is crucial for elucidating the effects of microplastics on cellular viability and for devising strategies to mitigate their potential adverse impacts.

### 3.2. Cell differentiation

Cell differentiation is paramount in the cultivated meat production process, serving as an indispensable determinant of the final product’s organoleptic and nutritional characteristics. This involves guiding pluripotent or multipotent cells, predominantly stem cells, through specific differentiation pathways to yield the requisite specialized cell types constituting meat, such as myocytes, adipocytes, and fibroblasts (Allan et al., 2019; Reiss et al., 2021; Zakrzewski et al., 2019). The meticulous orchestration of myocyte differentiation is pivotal for the formation of myofibrils, conferring the unique texture and mouthfeel characteristic of meat (K. Y. Lee et al., 2021; Listrat et al., 2016). In parallel, the directed differentiation of adipocytes is crucial for the deposition of intramuscular fat, a critical determinant of flavor profile and marbling (Li et al., 2020). Additionally, fibroblast differentiation and subsequent connective tissue formation offer essential structural integrity and have implications for meat tenderness (Purslow, 2020). Therefore, an in-depth understanding and precise control over these differentiation processes are indispensable for the optimization and scalability of cultivated meat, ensuring both its commercial viability and alignment with consumer expectations (Bomkamp et al., 2023; Reiss et al., 2021).

#### 3.2.1. Differential gene expression

In this study, we analyzed mackerel cells for the expression of three cardinal muscle differentiation markers, namely MYOD1, MYOG, and TNNT3A. MYOD1 is characteristic of myogenic progenitors (myoblasts) during the early phases of muscle cell differentiation, MYOG is expressed at the myocyte stage, and TNNT3A acts as a late marker associated with skeletal muscle function. Figure 3A illustrates the absolute gene expression values for MYOD1, MYOG, and TNNT3A of mack-2 cells cultivated in varying concentrations of microplastics (0, 1, 10, and 50 μg/mL). Remarkably, a heightened expression of MYOG compared to MYOD1 was observed, denoting that the cells were in a well-differentiated state. However, the absence of TNNT3A expression signified that the cells had not progressed to the later stages of muscle differentiation. Figures 3B and 3C depict the fold-change expression of MYOD1 and MYOG genes, represented as 2^(-ΔΔCt), over the control treatment with no microplastics. HPRT served as the housekeeping gene for expression normalization. Notably, a general lack of statistical difference was observed between the treatments in gene expression under varied microplastic concentrations, as calculated by one-way ANOVA. The exception was a single noteworthy difference in MYOD1 expression between 10 μg/mL and 50 μg/mL concentrations; however, this observation was tenuously supported with a p-value bordering on 0.05, indicating a weak statistical significance.

**Figure 3.**
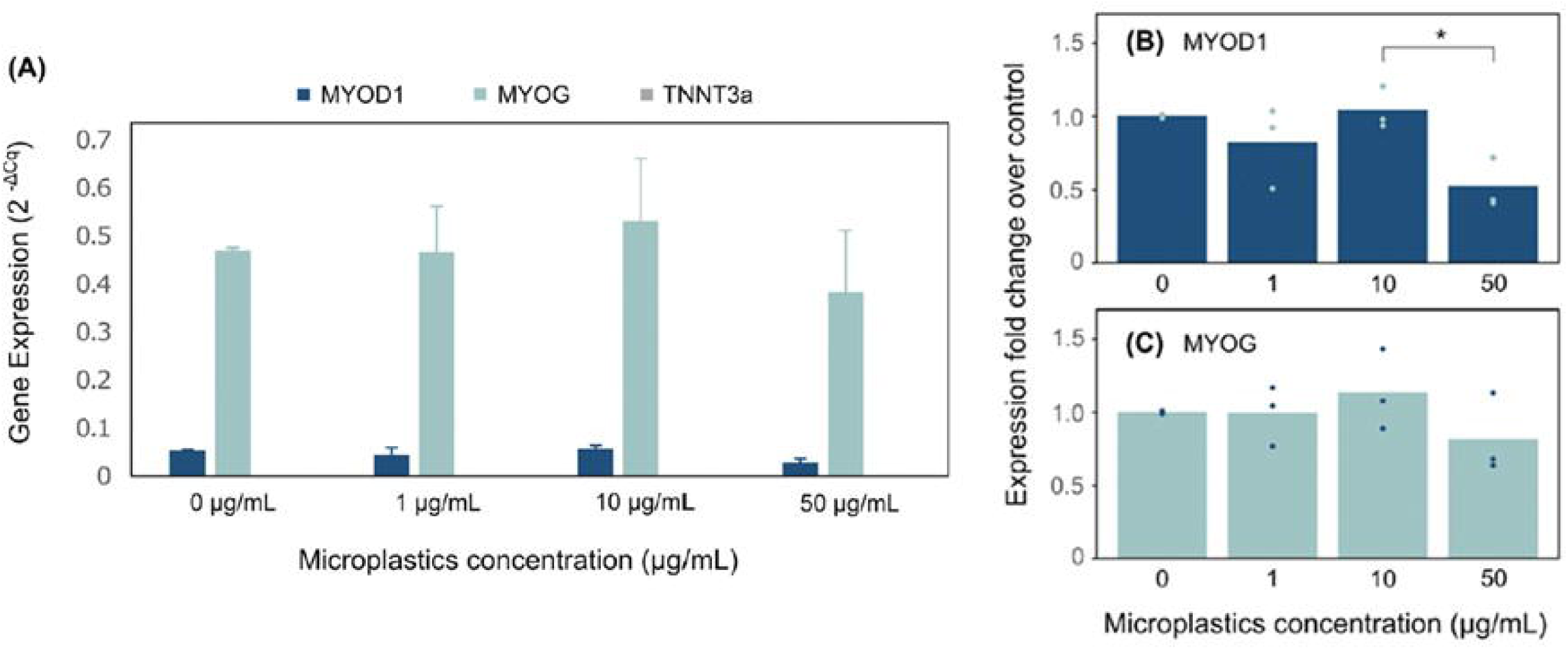
Expression of muscle differentiation gene markers in MACK cell growing with/without MPs: (A) Absolute gene expression values for MYOD1, MYOG and TNNT3A of MACK cells. Gene expression is represented as 2^(-ΔCt). (B, C) Fold-change expression of MYOD1 and MYOG genes over the control treatment. Points indicate single data points for each biological replicate and bars indicate their mean value (n = 3 experimental, n = 3 technical). HPRT was used as the housekeeping gene for expression normalization. Statistical significance calculated by one-way ANOVA is indicated by asterisks, in which p < 0.05 (*).

The observed increase in MYOG expression, coupled with the non-expression of TNNT3A, indicated a state of well-differentiated cells that had not yet reached the later stages of muscle development. This observation could be attributed to the potential onset of cellular senescence or reduced cell viability, especially considering that the mackerel cell line utilized was at passage #82, a stage at which cells often exhibit altered differentiation patterns due to accumulated genetic and epigenetic changes (Di Micco et al., 2021). Despite the variations in microplastic concentrations, the overall gene expression exhibited minimal statistically significant differences, emphasizing the need for further exploration into the interactions between microplastics and cell differentiation processes.

#### 3.2.2. Immunocytochemical analysis

Complementary to our qPCR findings, immunocytochemical assays provided a visual confirmation of cell differentiation processes. Cells subjected to various microplastic (MP) concentrations (1, 10, 50 ug/mL) over a 14 – day period, and a microplastic-free control were subsequently probed with MF20, DAPI, and Phalloidin, targeting myosin, cellular nuclei, and actin filaments, respectively. As depicted in Figure 4, each experimental condition displayed characteristic staining patterns across the trio of molecular markers. Notably, elongated structures positive for myosin, specifically stained by MF20, were evident in all conditions. This consistent staining pattern underscores mackerel cell differentiation into muscle cells. Importantly, the similarities observed between MP-exposed groups and the control suggest that the tested MP concentrations had a negligible impact on mackerel cell differentiation.

**Figure 4.**
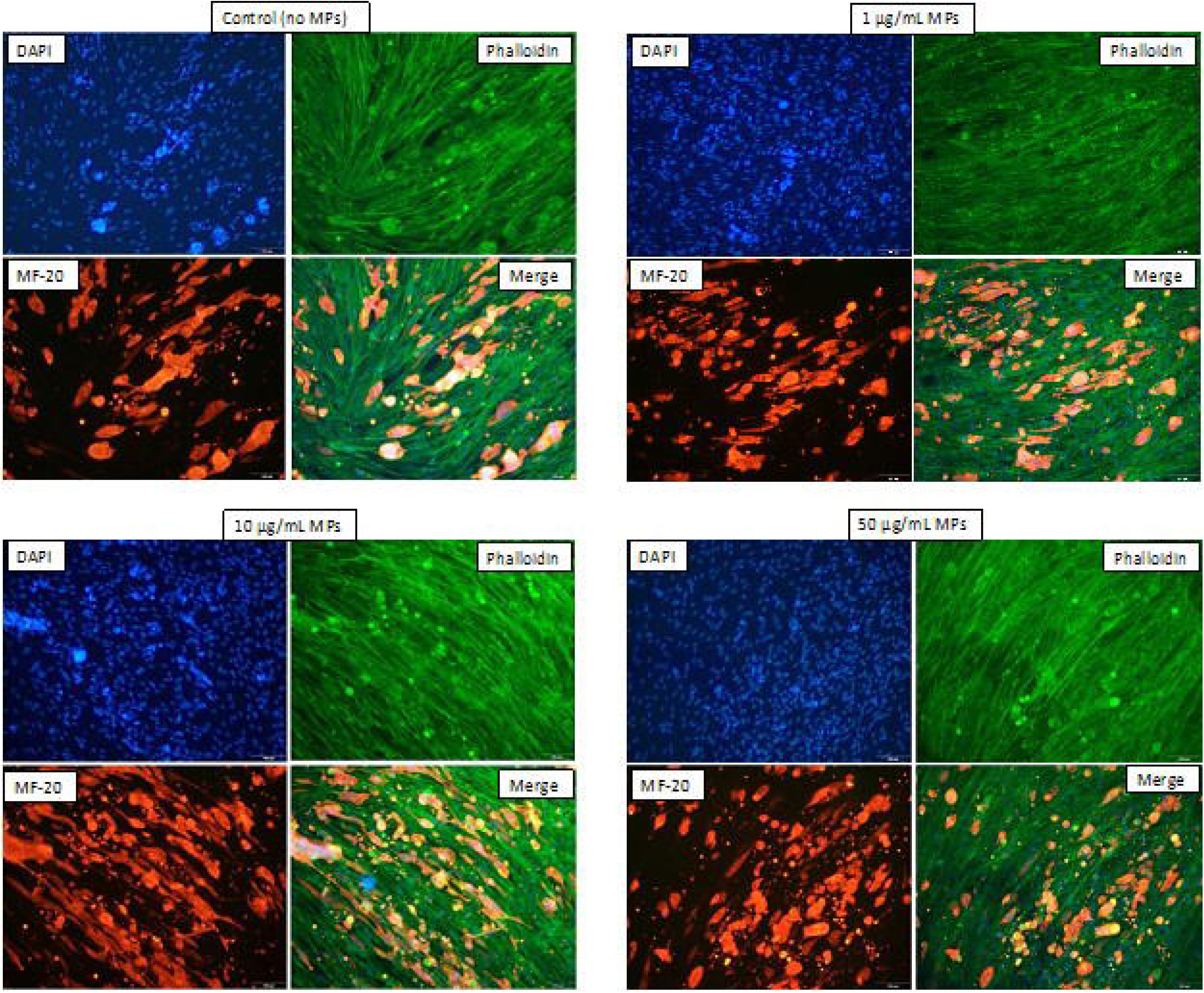
Representative immunostaining images. DAPI (blue) labels DNA, suitable for both live and fixed cells; Phalloidin (green) stains actin filaments in cells and tissues; MF20 (red), a monoclonal antibody, targets differentiated muscle fibers.

#### 3.2.3. Effects of microplastics on cell differentiation

While our findings demonstrated cell differentiation, the effects of microplastics on this process remained indiscernible; all treatment groups and controls yielded analogous outcomes. In contrast, earlier research has highlighted distinct impacts of microplastics on cell differentiation. For instance, Najahi et al. (2022) examined the influence of polyethylene terephthalate microplastics (MPs-PET, < 1 μm and < 2.6 μm) on human mesenchymal stromal cells, revealing a 30% reduction in cell proliferation and alterations in the differentiation potential of adipose and bone marrow cells. Concurrently, Han et al. (2020) found that polyvinyl chloride (PVC) and acrylonitrile butadiene styrene (ABS) microplastics influenced non-adhesive peripheral blood mononuclear cells (PBMCs) to differentiate into dendritic cells, implying that such plastic exposures might instigate human immune responses. In another study, Hua et al. (2022) highlighted that PS microplastics could disrupt cortical layer differentiation in cerebral spheroids, emphasizing the potential neurotoxic implications. Similarly, Im et al. (2022) reported that polystyrene nanoparticles, especially those with decreased crosslinking density, influenced reactive oxygen species activity and notably promoted adipogenic differentiation in mesenchymal stem cells. Moreover, it was reported that the stage of cell differentiation could influence the cell’s interaction with microplastics, especially regarding the uptake of these particles (Peng et al., 2023). In that study, researchers demonstrated that 2-μm polystyrene (PS) microplastics affected human cell lines differently based on their differentiation state, with undifferentiated Caco-2 cells showing significant PS uptake, whereas differentiated cells presented a reduced capacity for PS internalization. Collectively, these findings underscore the multifaceted impacts of micro/nanoplastics on cell differentiation across different cell types, emphasizing the need for comprehensive understanding and vigilant monitoring. Additionally, variables including the specific plastic type, exposure duration, and plastic concentration remain crucial determinants that could further modulate these effects (Smith et al., 2018).

## 4. Conclusion

In conclusion, this study evaluated the impact of microplastics on Atlantic mackerel (*S. scombrus*) skeletal muscle cell lines, using fluorescent polyethylene microspheres (10-45 µm) as model microplastics. The focus was primarily on understanding the effects of microplastic exposure on essential cellular processes, namely cell viability during attachment and growth phases and cell differentiation, which hold paramount significance in cultivated meat production. Utilizing microplastic concentrations of 1 µg/mL, 10 µg/mL, and 50 µg/mL, alongside a control devoid of microplastics, the study adopted the Trypan Blue Assay for cell viability assessment. The findings highlighted a marked difference in cell viability among microplastic-exposed treatments compared to the control. In parallel, cell differentiation was investigated using RT-qPCR for gene expression analysis and Immunostaining methodologies. Notwithstanding the observable cell differentiation, the study discerned no pronounced influence of microplastics on cell differentiation.

The findings of this study elucidate the interplay between microplastics and cellular mechanisms, highlighting the potential ramifications for cellular processes. As preliminary data, this investigation lays a foundation for subsequent, more detailed studies. Given the pervasive presence of microplastics in contemporary environments, it is imperative to explore their broader effects on cellular systems. Recognizing and understanding these implications is not only vital for advancing biotechnological applications but also for discerning potential long-term impacts on broader ecological and human health contexts.

## Acknowledgements

This research was financially supported by the Agriculture and Food Research Initiative (AFRI) Sustainable Agricultural Systems program, grant no. 2021-699012-35978 from the USDA National Institute of Food and Agriculture, and Texas A&M AgriLife Research. The authors would like to gratefully acknowledge Professor David Kaplan and Michael Saad for providing the cell line for this research.

## Notes

### Competing Interest Statement

The authors have declared no competing interest.

